# Fourier SPoC: A customised machine-learning analysis pipeline for auditory beat-based entrainment in the MEG

**DOI:** 10.1101/2021.04.23.441088

**Authors:** Stephanie Brandl, Niels Trusbak Haumann, Simjon Radloff, Sven Dähne, Leonardo Bonetti, Peter Vuust, Elvira Brattico, Manon Grube

## Abstract

We propose here (the informed use) of a customised, data-driven machine-learning pipeline to analyse magnetoencephalography (MEG) in a theoretical source space, with respect to the processing of a regular beat. This hypothesis- and data-driven analysis pipeline allows us to extract the maximally relevant components in MEG source-space, with respect to the oscillatory power in the frequency band of interest and, most importantly, the beat-related modulation of that power. Our pipeline combines Spatio-Spectral Decomposition as a first step to seek activity in the frequency band of interest (SSD, [1]) with a Source Power Co-modulation analysis (SPoC; [2]), which extracts those components that maximally entrain their activity with the given target function, that is here with the periodicity of the beat in the frequency domain (hence, f-SPoC). MEG data (102 magnetometers) from 28 participants passively listening to a 5-min long regular tone sequence with a 400 ms beat period (the “target function” for SPoC) were segmented into epochs of two beat periods each to guarantee a sufficiently long time window. As a comparison pipeline to SSD and f-SpoC, we carried out a state-of-the-art cluster-based permutation analysis (CBPA, [3]). The time-frequency analysis (TFA) of the extracted activity showed clear regular patterns of periodically occurring peaks and troughs across the alpha and beta band (8-20 Hz) in the f-SPoC but not in the CBPA results, and both the depth and the specificity of modulation to the beat frequency yielded a significant advantage. Future applications of this pipeline will address target the relevance to behaviour and inform analogous analyses in the EEG, in order to finally work toward addressing dysfunctions in beat-based timing and their consequences.

**Author summary:** When listening to a regular beat, oscillations in the brain have been shown to synchronise with the frequency of that given beat. This phenomenon is called entrainment and has in previous brain-imaging studies been shown in the form of one peak and trough per beat cycle in a range of frequency bands within 15-25 Hz (beta band). Using machine-learning techniques, we designed an analysis pipeline based on Source-Power Co-Modulation (SPoC) that enables us to extract spatial components in MEG recordings that show these synchronisation effects very clearly especially across 8-20 Hz. This approach requires no anatomical knowledge of the individual or even the average brain, it is purely data driven and can be applied in a hypothesis-driven fashion with respect to the “function” that we expect the brain to entrain with and the frequency band within which we expect to see this entrainment. We here apply our customised pipeline using “f-SPoC” to MEG recordings from 28 participants passively listening to a 5-min long tone sequence with a regular 2.5 Hz beat. In comparison to a cluster-based permutation analysis (CBPA) which finds sensors that show statistically significant power modulations across participants, our individually extracted f-SPoC components find a much stronger and clearer pattern of peaks and troughs within one beat cycle. In future work, this pipeline can be implemented to tackle more complex “target functions” like speech and music, and might pave the way toward rhythm-based rehabilitation strategies.

## 1 Introduction

Rhythm and timing are crucial in our daily lives. Sequencing information at the level of a few hundred milliseconds in particular plays a major role for both the perception and production of music, speech, and movement. This relevant time scale here corresponds to that of our ability to “feel the beat”, that is from approximately 200 to 800 ms [4]. The ability to detect and process a regular rhythmic beat in an auditory stream of acoustic events is one of the essential mechanisms to our musical, speech and language skills (e.g., [5]). This ability can be disturbed in neurological as well as developmental disorders with marked consequences to the individuals’ life quality, in terms of the perception and or production of music, speech, or bodily movement (e.g. [6–8]). It is therefore of critical importance to first understand how the brain tracks the beat, to then find ways to monitor and support the rehabilitation of dysfunctional timing processes in individuals with those disorders.

The human brain’s timing system comprises a number of structures that were traditionally associated with motor functions and include the basal ganglia, cerebellum, and parietal, temporal and frontal areas. The activation of this network of structures has been indicated by neuroimaging work [9–11], and corroborated by neuropsychological [6, 12] and transcranial magnetic stimulation studies [13], two essential methodologies that support a direct test of the criticalness of the given brain structures. Each of the timing network’s components has to a different degree been associated with specific aspects of motor or sensory timing, or beat-vs. duration-based, implicit or explicit timing. The supplementary motor area and basal ganglia for instance have been repeatedly reported to be more engaged in beat-based timing [14], and that with further sub-specialization in finding vs. keeping the beat for instance [15]. However, findings are not entirely consistent in terms of the specificity of criticalness (e.g. [16, 17]), in part probably due to differences in lesions or tasks. The most important point here might be, that the behaviourally measurable outcome as well as the neural processes will in most instances be a network result [18, 19] rather than that of a one structure alone. In terms of neural mechanisms, one that has been emerging from recent literature is that of coupled oscillations, i.e. the entrainment of the power of a faster oscillation with the phase of the slower (e.g. [20]). More specifically, the timing system has been promoted to feature the entrainment of neural oscillations in the beta range (13-30 Hz) with the periodicity of the beat in musical rhythms [21].

In attempts to get to the bottom of the structural and functional neural correlates that underlie our entrainment with and “feeling of the beat”, previous MEG and EEG work both demonstrated clear effects in oscillatory activity. In their seminal 2012 MEG study, Fujioka et al. [22] demonstrated a clear periodic modulation in the beta band power, with the beat frequency in the MEG. The data showed one peak and one trough per beat cycle within the beta band and the same pattern could be seen for an isochronous (i.e. perfectly regular) beat for three different tempi of 390, 585, and 780 ms. These activity patterns were extracted as the result of an anatomically guided analysis and appeared most prominently in the auditory cortex of the temporal lobe. Further support for the beta band “driving the brain on beats” came from related EEG work ([23], Chang / Tillmann ?) with converging conclusions [24].

We here present an alternative, customised, and open-source machine-learning analysis pipeline that enables us to extract in a data-driven way the maximally relevant brain activity in source space. The advantages of this pipeline are that no a priori input about brain structures is required, neither in terms of pre-defined anatomical brain regions of interest nor in terms of the individual’s brain. In fact, this pipeline may not only pick up on the activity of individual brain structures, but also the timing network activity or the communication between its components that may happen in a band-specific fashion of the timing network than that of individual structures. Our pipeline entails two spatial filtering techniques: Source Power Co-modulation Analysis (SPoC, [2]), and Spatio-Spectral Decomposition (SSD, [1]). SPoC, which lies at the heart of this pipeline, allows us to seek components in source space that maximally vary their activity with a function of interest, i.e. the periodicity of the beat (here, 400ms). The application of SSD prior to SPoC optimizes the input for SPoC with respect to the activity in the band of interest, initially. In addition to the pre-defined frequency band of interest in the beta range, we sought components that might vary their power in the alpha and gamma band, for which results in the pre-existing literature were much less prominent [21, 25]. In order to test the efficacy of our analysis pipeline, we employ a most sophisticated sensor-space analysis, based on a state-of-the-art cluster-based permutation analysis (CBPA; [3]). To compare the results between the two, we employ a wavelet-based time-frequency analysis (TFA) followed by a fast-Fourier transformation (FFT) to calculate the depth and specificity of the periodic modulation of activity as two quantifiable outcome measures. The final goal of this analysis is to provide a data-driven assumption-free brain marker of neuronal entrainment to an external beat.

Code for this pipeline will be made publicly available on GitHub.

## 2 Methods

In order to investigate on neural oscillations that are expected to entrain to a regular auditory input [22], we recorded the MEG of participants hearing a metronome-like steady repetition of a tone.

### 2.1 MEG Recordings

The magnetoencephalography (MEG) data was acquired by employing an Elekta Neuromag TRIUX system (Elekta Neuromag, Helsinki, Finland) equipped with 306 channels. The machine was positioned in a magnetically shielded room at Aarhus University Hospital, Denmark. Data was recorded at a sampling rate of 1000 Hz with an analogue filtering of 0.1–330 Hz. Prior to the measurements, we accommodated the sound volume at 50 dB above the minimum hearing threshold of each participant. Moreover, by utilizing a 3D digitizer (Polhemus Fastrak, Colchester, VT, USA) and a continuous head position identification (cHPI) system, we registered the position over time of 4 headcoils, with respect to 3 anatomical landmarks (nasion, and left and right preauricular locations). This procedure allowed us to track the exact head location within the MEG scanner at each time-point. We utilized this data to perform an accurate movement correction at a later stage of data analysis. The MEG was recorded while participants performed a passive listening task while watching a silent movie. The auditory stimuli consisted of a continuous sequence of 300 Hz sine tones of duration 100 ms (with 10 ms fade-in and -out) presented with an isochronous inter-onset interval of 400 ms for a total duration of 5 min. The present study was conducted on a subset of a larger dataset acquired at Aarhus University and approved by the Ethics Committee of the Central Denmark Region (De Videnskabsetiske Komiteer for Region Midtjylland) (ref. 1-10-72-411-17). Additionally, the whole procedure was carried out according to the Declaration of Helsinki [26].

### 2.2 Participants

We recruited 28 participants whose brain activity was recorded using MEG, 18 female (62%) and 11 male (38%). The age of the participants ranged from 19 to 50 years (m = 26.45, SD = 7.37). The group consisted of 4 first-year music students and 25 non-musicians. The latter reported to have no more than 2 years of formal musical theory training (with the exception of one participant reporting 4-6 years, m = 0.23, SD = 0.47) and no more than 5 years of formal musical instrument training (m = 0.59, SD = 1.10). All participants declared to be in good health, and not to suffer from any neurological disease. Additionally, no psychiatric medication was reported and no hearing impairment was observed. The participants were all right-handed with one exception being left-handed. The participants originated from 14 different countries and 7 of them (24%) were Danish which was the largest national group. We compensated the participants for the time spent in the experiment with vouchers for online-shops. On arrival, participants were briefly instructed about the experimental procedures prior to sign the informed consent and begin the experiment. A researcher was present at any time for assistance and participants were given the possibility to withdraw their participation in the study at any time.

### 2.3 Preprocessing

The raw MEG sensor data (204 planar gradiometers and 102 magnetometers) was pre-processed by the Elekta Neuromag MaxFilter(TM) software version 2.2.15 [27] [28] for attenuating the interference originated outside the scalp by applying temporal signal space separation (tSSS). Within the same session, Maxfilter also adjusted the signal for head movement. The default tSSS parameters were applied with inside expansion order of 8, outside expansion order of 3, optimization of both inside and outside bases, default subspace correlation limit of 0.980, default raw data buffer length of 10 seconds. Subsequent data preprocessing steps were performed by using the FieldTrip Toolbox (2018-11-12) for Matlab [29] on the broadly distributed magnetic fields from the 102 axial magnetometer sensors only to allow a better comparison with EEG and other MEG systems. Therefore, the narrow gradients of the 204 planar gradiometers were not considered here. Data has been low- and high-pass filtered at 500 Hz and 0.5 Hz, respectively, with a finite impulse response filter. To remove artifacts introduced by heart beat and eye movements, independent component analysis has been applied [30] and up to 2 out of 64 components per participant (one for heart beat and one for eye movements) have been removed after visual inspection. We then cut the continuous data into epochs of two beat periods (800ms + 1000ms before and after as a buffer for the wavelet analysis) and aligned them 1000ms prior to tone onset.

Finally, we split the data into train and test set, we therefore took out every third tone and defined this as the test set which we excluded from the analysis steps 3.1.1 and 4.1.1-4.1.3. This is standard protocol in most machine learning-based approaches where an algorithm needs to be fitted on seen data, for instance when hyperparameters need to be set. To evaluate if the “trained” algorithm also works on new/unseen data it is then applied to data it has not been trained/fitted on, this way we prevent the algorithm from overfitting.

## 3 Sensor-Level Analysis

As a first approach to detect beat-based synchronization/desynchronization effects and as a comparison for SPoC results, we wanted to investigate data-driven methods at sensor-level. We therefore conducted cluster-based permutation analysis to identify significant sensors from which we could later compute the time-frequency response. This serves as a baseline to our machine learning-based spatial filtering approach (see Sec. 4).

### 3.1 Analysis Pipeline

#### 3.1.1 Cluster-based permutation analysis

We applied a one-sample cluster-based permutation test for MEG [3], which finds statistically significant power modulations across participants. This is a data-driven statistical method that is useful in finding statistically significant clusters of power synchronization/desynchronization spanning coherent points in time, frequency, and sensors location where the relevant points in time, frequency and sensor location are unknown. The permutation version of the one-sample t-test iteratively performs one-sample t-tests with random signs added to the observed sample to estimate a null-distribution of t-values. The proportion of t-values in this null-distribution exceeding the observed t-value (obtained without adding random signs) indicates the p-value [31]. It should be noted that for the permutation tests, including the one-sample version, the data can be non-normally distributed, however, for the one-sample test it is assumed that the population values are symmetrically distributed [31], which is a reasonable assumption given that the power modulations are periodic. The cluster-based version of the permutation test includes an initial step, where neighboring significant points in time, frequency and sensor location are merged into larger clusters, thereby offering a solution to the need for multiple tests conducted on each point that is typically not independent in MEG [3]. For the cluster-based permutation test we carried out 1000 randomizations. For the initial cluster statistic we used the default channel neighbor configuration for the Neuromag Triux MEG system implemented in FieldTrip, the maximum sum t-statistic, and a significance threshold of p=0.01. For the cluster-based permutation statistics we used a significance threshold of p=0.025.

We considered all 102 axial magnetometer sensors, 20 logarithmically scaled frequency bins between 5-35 Hz and 100 linearly scaled time points in a time window of 400ms corresponding to one beat period. This resulted in findings of 3 desynchronization [negative] and 2 synchronization [positive] clusters in time, frequency, and sensor location that were significant across participants.

We further found 78 sensors that proved to be significant across all participants in the cluster-based permutation analysis. For each sensor we therefore averaged over a 2-dimensional binary array of all frequency bins and time points (29 × 100) where 1 means that the corresponding sensor at an individual time point and frequency bin is significant. Each sensor with an average value larger than 0, i.e. each sensor that is significant in at least one (frequency, time) combination, is considered as significant. The average values, i.e. the average percentage of significance for each sensor is displayed in Fig. 1.

**Fig 1.**
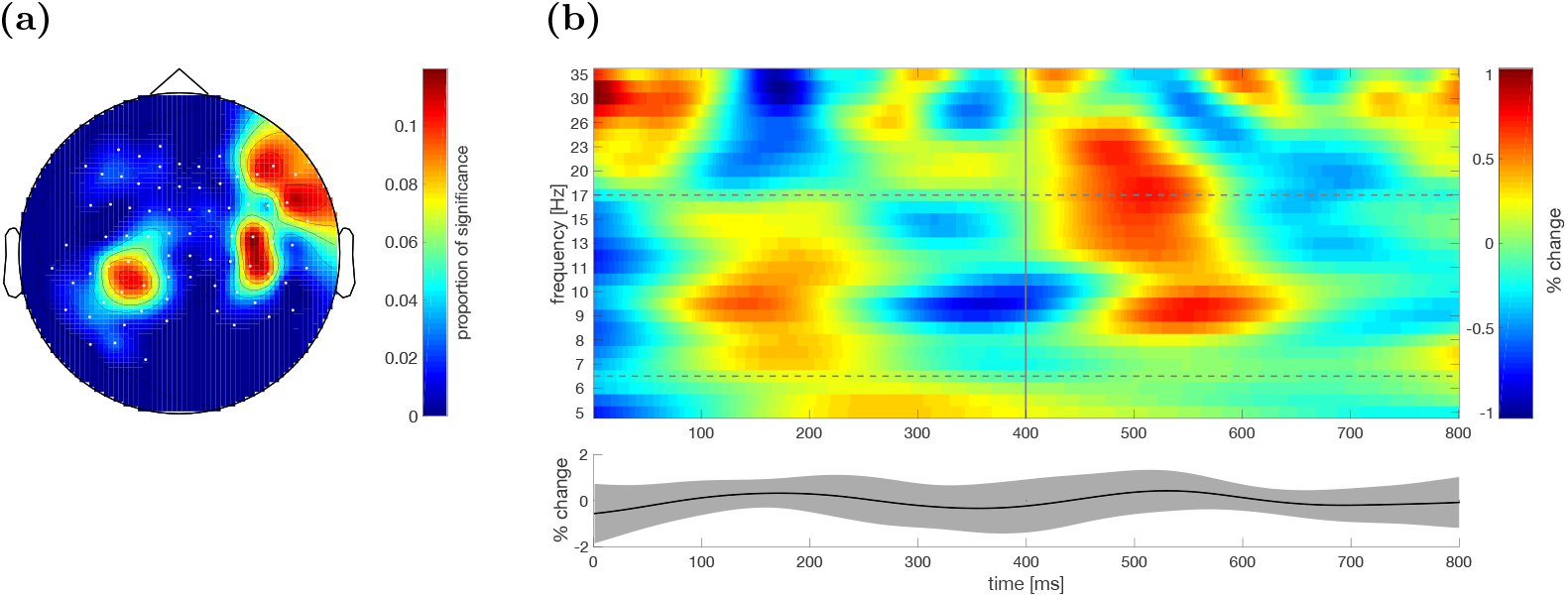
Results of cluster-based permutation analysis (CBPA) *(a)* The topography shows the average proportion of significance for each sensor averaged over participants and frequency bins. The 78 sensors that proved to be significant across all participants in the CBPA are highlighted by a white dot. *(b)* Time frequency response (TFR, *upper*) averaged across participants, time windows and significant sensors shows percentage change in power. In the *lower* subplot averaged across the frequency band of 6-18 Hz TFR shows most regular activity with a peak around 150ms after tone onset and a through around 300ms after tone onset. The grey area shows standard deviation across participants.

#### 3.1.2 Spectral Analysis

Time-frequency analysis (TFA) was applied to the 78 sensors that proved to be significant in the cluster-based permutation analysis [32]. We applied wavelet analysis to individual time windows (onset-aligned) of 2-beat periods for all 78 significant sensors in a frequency band of 5-35 Hz, individually for each participant (Morlet wavelet, window width of 7). We computed the spectral amplitude (after removing the buffer window), averaged over trials and detrended. We then calculated the percentage change of power by subtracting and dividing by the mean of the entire trial length for each frequency bin individually. Finally, we averaged over sensors and participants.

### 3.2 Results

In Fig. 1 we show results of the cluster-based permutation analysis and the corresponding time frequency response (TFR). The topography on the left shows the average percentage of significance for each sensor. The significant clusters of power changes were overlapping in channel, time and frequency extended between 4 to 360 ms after the beat stimulus onset and between frequencies 6-30 Hz.

On the upper right we visualized the time frequency response in 5-35 Hz averaged over all 78 significant sensors and participants. We see a clear regular event-related synchronisation/ desynchronisation (ERS/ERD, [33]) in the frequency band of 6-18 Hz with a synchronisation starting around 50 ms after each stimulus onset and a desynchronisation starting 280 ms after stimulus onset up to 50 ms after the next stimulus onset with up to ±1% power change.

On the lower right we further averaged over the frequency band of 6-18 Hz, i.e. an average envelope where we see a regular ERS/ERD even clearer with a positive peak around 150 ms after stimulus onset and a negative peak around 350 ms after stimulus onset.

## 4 Spatial Filtering

To extract an amplitude-modulated oscillatory component, we applied spatial filtering methods to the preprocessed MEG data. Spatio-Spectral Decomposition computes filters that improve signal-to-noise-ratio in a frequency band of interest which here serves as a preprocessing step to find a lower dimensional representation, we afterwards apply a novel version of Source Power Co-Modulation to extract components that maximally correlate with the modulation of the beat. It is important to note that we do not need any anatomical knowledge of the individual brain to extract those components. Both methods are purely mathematical and based on inverting the forward model [34]. This makes it easy to apply them to multidimensional brain signals such as EEG or MEG.

### 4.1 Analysis Pipeline

#### 4.1.1 Spatio-Spectral Decomposition

Spatio-Spectral Decomposition (SSD, [1]) extracts oscillatory signals with a high signal-to-noise-ratio (SNR). Spectral power in the frequency band of interest (inner SSD band) is being maximized while power in left and right neighboring frequency bands (outer SSD band) is minimized:

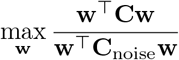

Here, **C** denotes the covariance matrix of the signal in the frequency band of interest, whereas **C**_noise_ denotes the covariance matrix of the signal in the neighboring frequency bands. Spatial filters are displayed as **w**.

This optimization problem can be solved with a generalized eigenvalue decomposition, more details can be found in [1].

We select the first 15 SSD filters for each participant, i.e. the filters that correspond to the 15 largest eigenvalues. Data from the 102 axial magnetometer sensors (see Sec. 2.3) is then projected to a 15-dimensional SSD-subspace, i.e.

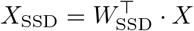

where 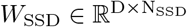 is the matrix with 15 SSD filters, *X* ∈ R^D×N^ the measured MEG data, 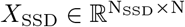 the projected MEG signal. *D* refers to the number of sensors, *N*_*SSD*_ to the number of selected SSD filters and *N* refers to the number of measured time points (see Table 1).

**Table 1.** Notation of parameters used in the analysis pipeline.

| Parameter | Meaning | Values |
| --- | --- | --- |
| N | Number of measured time points |  |
| D | Number of MEG sensors | 102 |
| $N_{SSD}$ | Number of selected SSD filters | 15 |
| $N_{SPoC}$ | Number of selected SPoC filters | 1 |

#### 4.1.2 Source Power Co-Modulation

Source Power Co-Modulation (SPoC_λ_, [2]) computes a set of spatial filters **W** which directly optimize the co-modulation between a target variable *z* and the power time course of a spatially filtered signal Φ over epochs *e*.

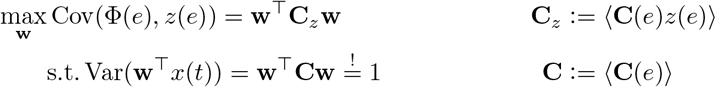

where ⟨ ⟩denotes the mean over epochs *e* and **C**(*e*) represents the covariance matrix of the e-th epoch.

This can then be solved by a generalized eigenvalue problem

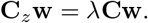

where the eigenvalue λ can be interpreted as the covariance between Φ and *z*. For further details, we refer the interested reader to the authors of SPoC, Dähne et al., [2].

For the special case of a regular target function, we propose Fourier-SPoC, an extension to SPoC_λ_.

#### 4.1.3 Fourier-SPoC

Since we are interested in finding regular synchronisation and desynchronisation effects within a specific frequency band we need to choose a regular target function for the optimization of the SPoC_λ_-filters. We therefore choose a sine function as the target function *z*(*e*) for fourier-SPoC (f-SPoC) as displayed in the lower part of Fig. 2. We thus optimize f-SPoC over phases of a sine function modulated here with the beat frequency of 2.5 Hz as we assume a rhythmic entrainment with the beat frequency. First, we bandpass-filter the SSD-transformed data with a passband in the range of the inner SSD band with a 3rd order zero-phase Butterworth filter. We then apply f-SPoC to the bandpass-filtered data and optimize over a grid of 10 different phases individually for each participant. Since we are looking for non-phase locked activity we do not know in which phase of the sine function the entrainment is happening. This means, we compute a set of SPoC filters for each of the 10 phases and select the sine function that maximally correlates with the envelope of the SPoC-filtered signal. We select the first and second SPoC filter for the respective phase, i.e. the ones with the largest eigenvalue and project the SSD-filtered signal *X*_SSD_ to this two-dimensional SPoC subspace.

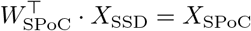

where *W*_SPoC_ ∈ℝ^15×2^ contains the first and second SPoC filter, *X*_SSD_ the SSD transformed MEG signal as described in Section 4.1.1 and *X*_SPoC_ ℝ^1×N^ the projected signal.

**Fig 2.**
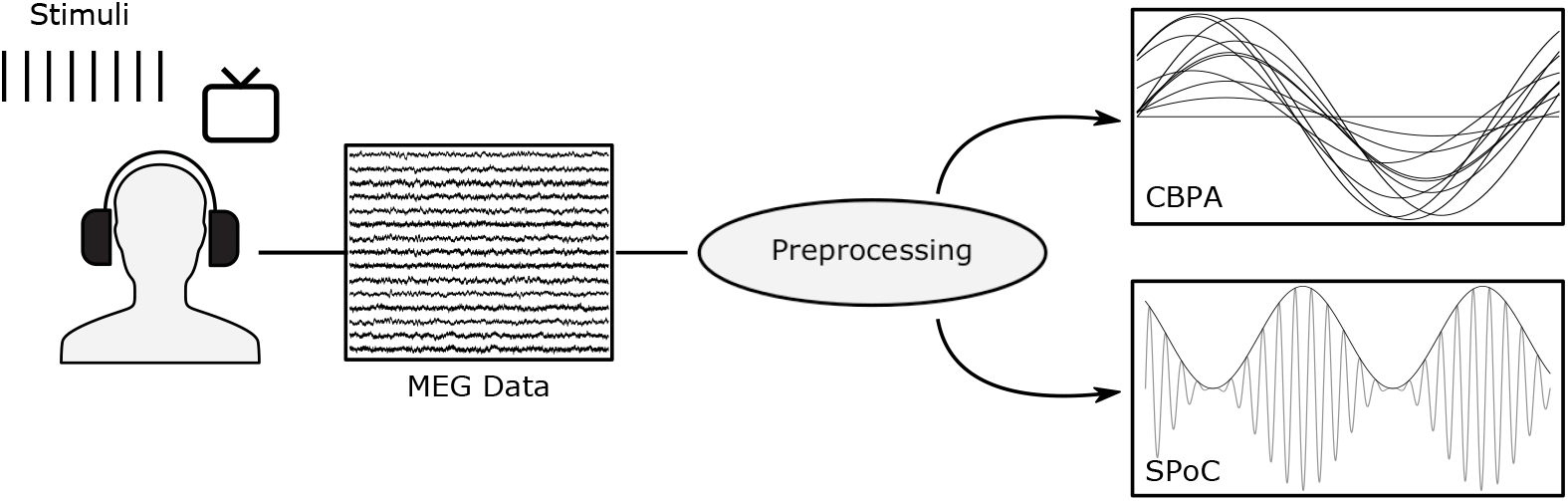
Overview over the two analysis approaches. From left to right: MEG data of participants passively listening to a regular beat were recorded while they were watching a silent movie. Data was then preprocessed (high- and lowpass filtering, ICA and cutting into windows of 2 beat periods) and we applied cluster based permutation analysis and a spatial filtering method (SPoC) separately.

Spectral analysis is then applied to the SPoC-filtered time course *X*_SPoC_ as described in Section 3.1.2 with the difference that we do not average over sensors but look at the two SPoC components individually.

### 4.2 Results

We applied the analysis pipeline (Sec. 4.1) to the recorded MEG data of each participant individually. Based on previous work, we selected three different frequency bands in the range of the alpha and beta band (which have been found to show entrainment): alpha (8-12 Hz), low beta (15-20 Hz) [23, 25, 35] and high beta (20-25 Hz) [35]. This means, we feed the respective frequency band as the frequency band of interest (inner SSD band) into the SSD algorithm and apply f-SPoC afterwards.

In Fig. 3 we show the time frequency responses (TFR) for the first and second SPoC-components for each frequency band. For all 3 frequency bands we show TFR in the range of 5-35 Hz and below an average envelope over the band of interest.

**Fig 3.**
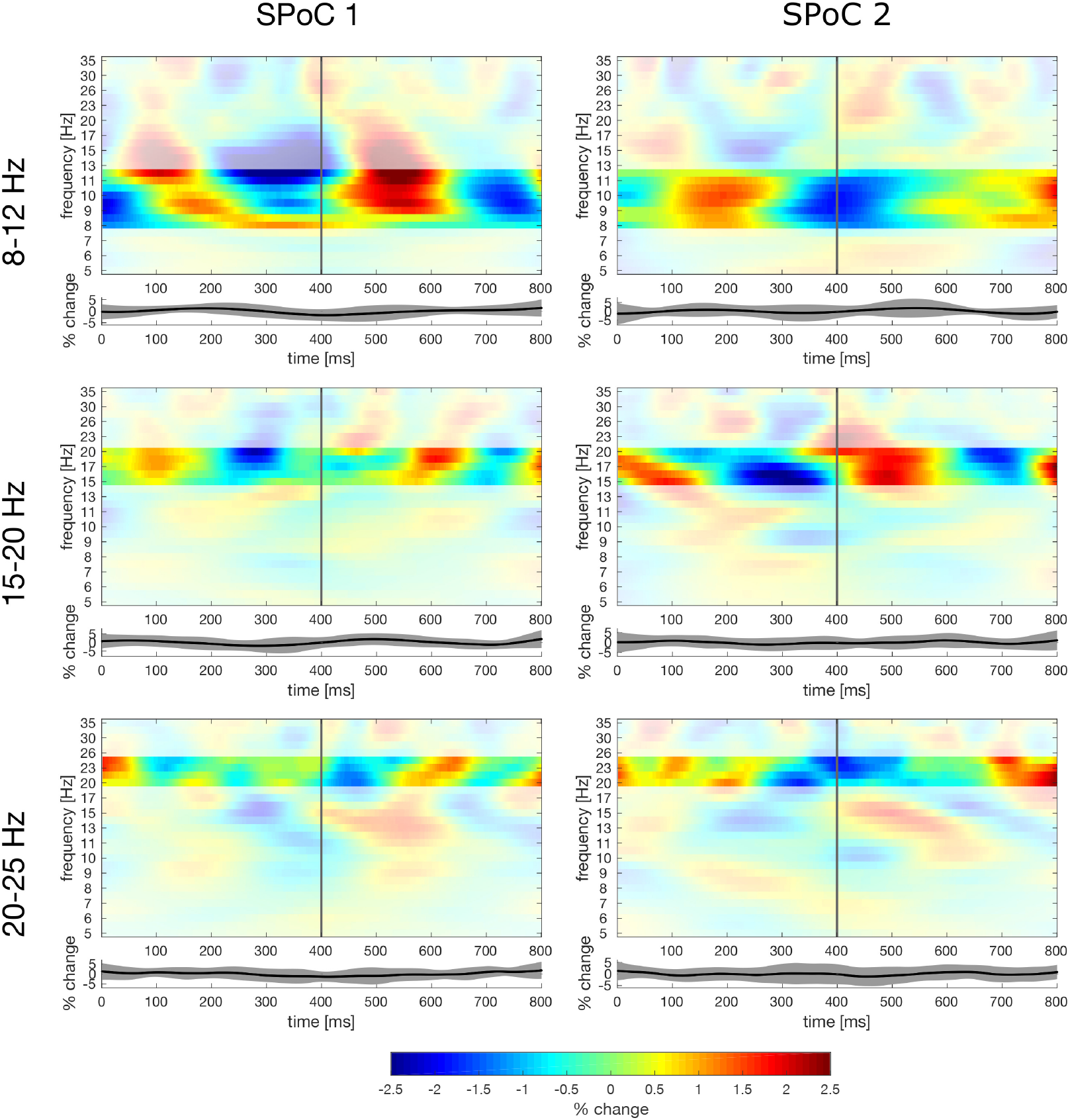
Time frequency response (TFR) of first and second SPoC component shows percentage change in power for different frequency bands averaged over all participants. Upper plots show TFR for 5-35 Hz with band of interest highlighted, lower plots show the average envelope over band of interest with the standard deviation across participants displayed as a grey area around the solid black line. Periodic peaks and troughs are most visible in the range of 8-20 Hz (upper left, center right and last row).

Clearest ERS/ERD effects are visible in the alpha band. Here, we see a clear synchronisation in the first SPoC component especially in the frequency band of 9-17 Hz with a peak around 150 ms after stimulus onset followed by a clear desynchronisation about 350 ms after stimulus onset up to 20 ms after the next stimulus with up to ±3% power change. The envelope below the TFR shows an average over the frequency band of 8-12 Hz over all participants. Also here we see the clear rhythmic ERS/ERD effects after stimulus onset.

For low beta (15-20 Hz) we see a similar pattern with a synchronisation around 100 ms after stimulus onset and a desynchronisation around 300 ms after stimulus onset with up to ±2.5% power change in the second component.

For high beta (20-25 Hz, lower left) the patterns are not clearly visible within 20-25 Hz but are still prominent in 10-17 Hz. Since the results for the alpha band show a clear ERS/ERD ranging further than the band of interest, we combine frequency bands of alpha and low beta.

It is important to note that in the SSD the power in the outer frequency band is tried to be minimised while power in the band of interest is increased. This means ERS/ERD effects present in both, the frequency band of interest and the neighboring frequencies might not be as clear in the resulting signal. Also, SPoC is based on a generalized eigenvalue decomposition which is known to be prone to outliers. This means that relevant information can also be captured by the second instead of the first component/eigenvector which we can see here for low beta.

The resulting TFR for the combined frequency band of alpha and low beta in Fig. 5 shows very clear synchronization and desynchronization effects in the frequency range of 8-20 Hz with up to ±5% power change. The envelope below shows an even more consistent and regular course than before in Fig. 3.

In Fig. 4, we show individual results of three exemplary participants for SPoC optimised over 8-20 Hz and CBPA as shown as group averages in Fig. 1 and Fig. 5. Individuals were handpicked to show the range of results on the individual level. SPoC and CBPA for the first participant both show periodic patterns while the former shows a much clearer and stronger pattern than the sensor-level analysis. SPoC analysis for the second participant still shows a very clear pattern while CBPA results in a rather unclear pattern. SPoC and CBPA for the third participant both do not show periodic patterns.

**Fig 4.**
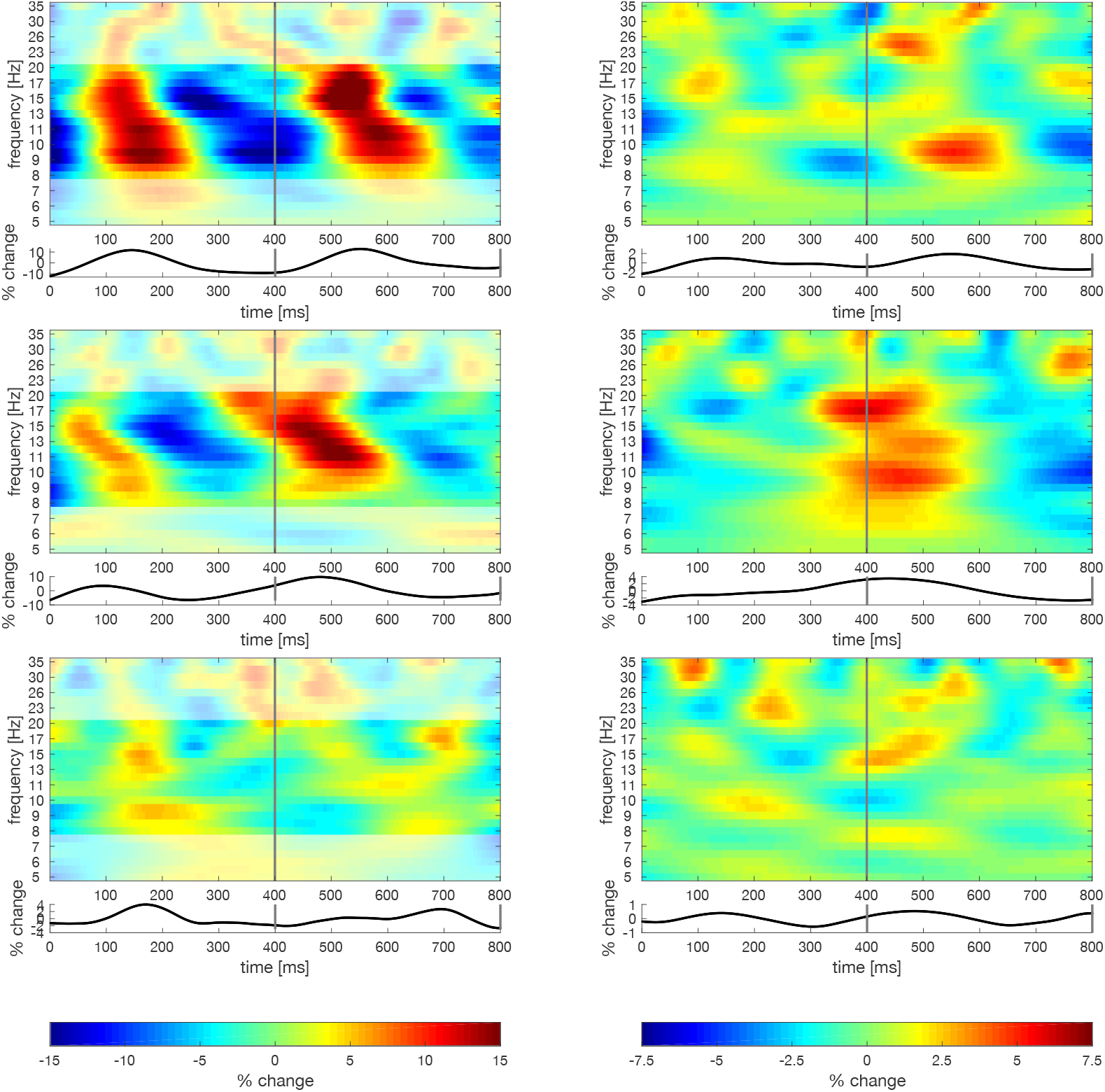
Time frequency response (TFR) of first SPoC component (left) and CBPA (right) shows percentage change in power for three individual participants (one per row) in the range of −15/15 % and −7.5/7.5 % for SPoC and CBPA respectively. Upper plots show TFR for 5-35 Hz with band of interest (8-20 Hz) highlighted in SPoC, lower plots show the average envelope over 8-20 Hz (SPoC) and 8-16 Hz (CBPA).

**Fig 5.**
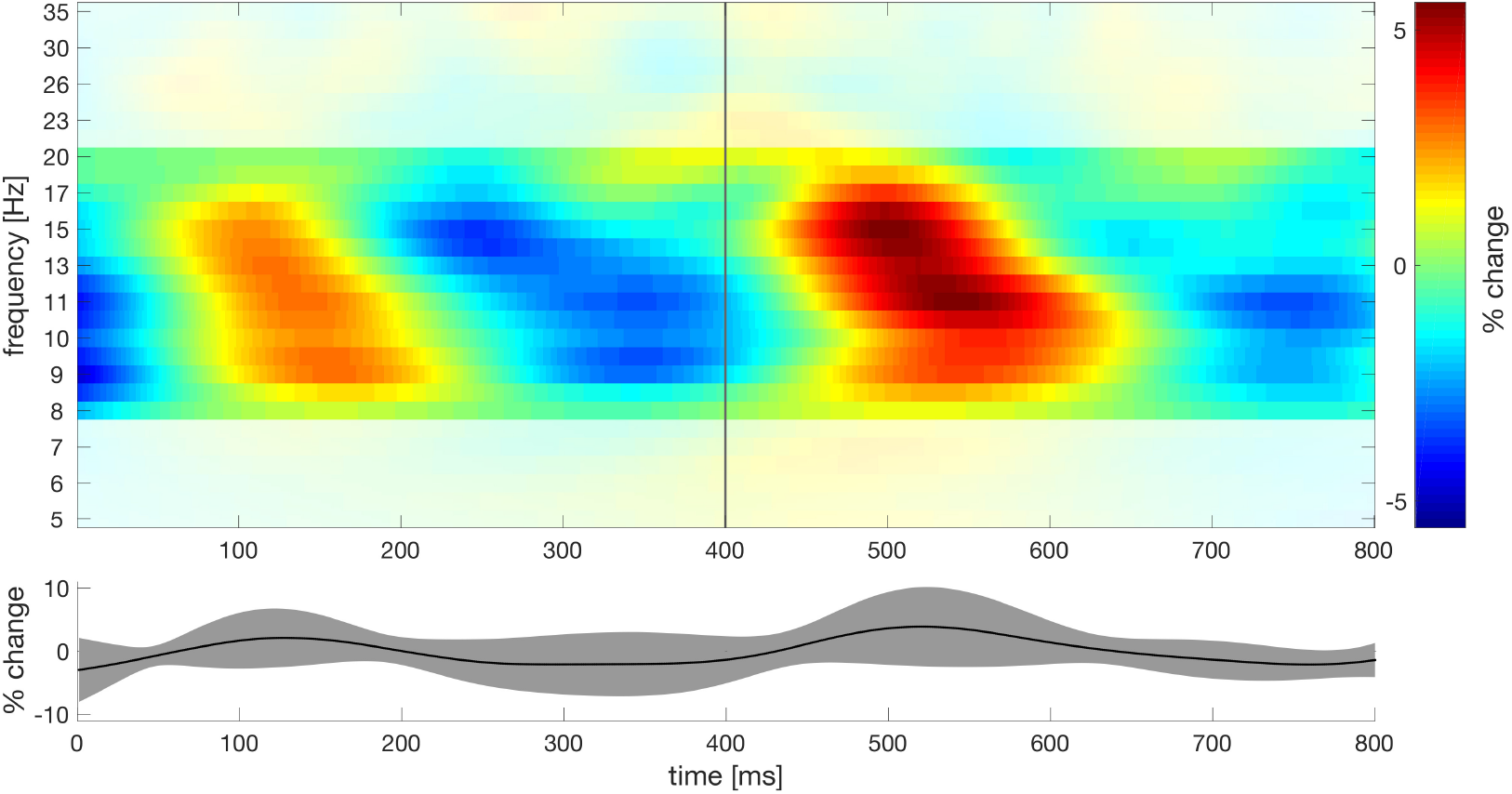
Time frequency response (TFR) of first SPoC component for the frequency band of 8-20 Hz. Lower plot shows the average envelope over the band of interest. Changes of up to 10% in power are visible after beat onset at 120 and 520 ms.

We quantify event-related synchronisation and desynchronisation for each participant by the difference between the maximal positive amplitude and the maximal negative amplitude in the average envelope of 8-20 Hz (SPoC) and 6-18 Hz (78 significant sensors in CBPA). Values for each participant are plotted in Fig. 6 (*left*). A higher value here means more power across the respective frequency band over the average epoch. Results for SPoC are significantly higher (*p <* 10^*−*4^) than the ones for CBPA.

**Fig 6.**
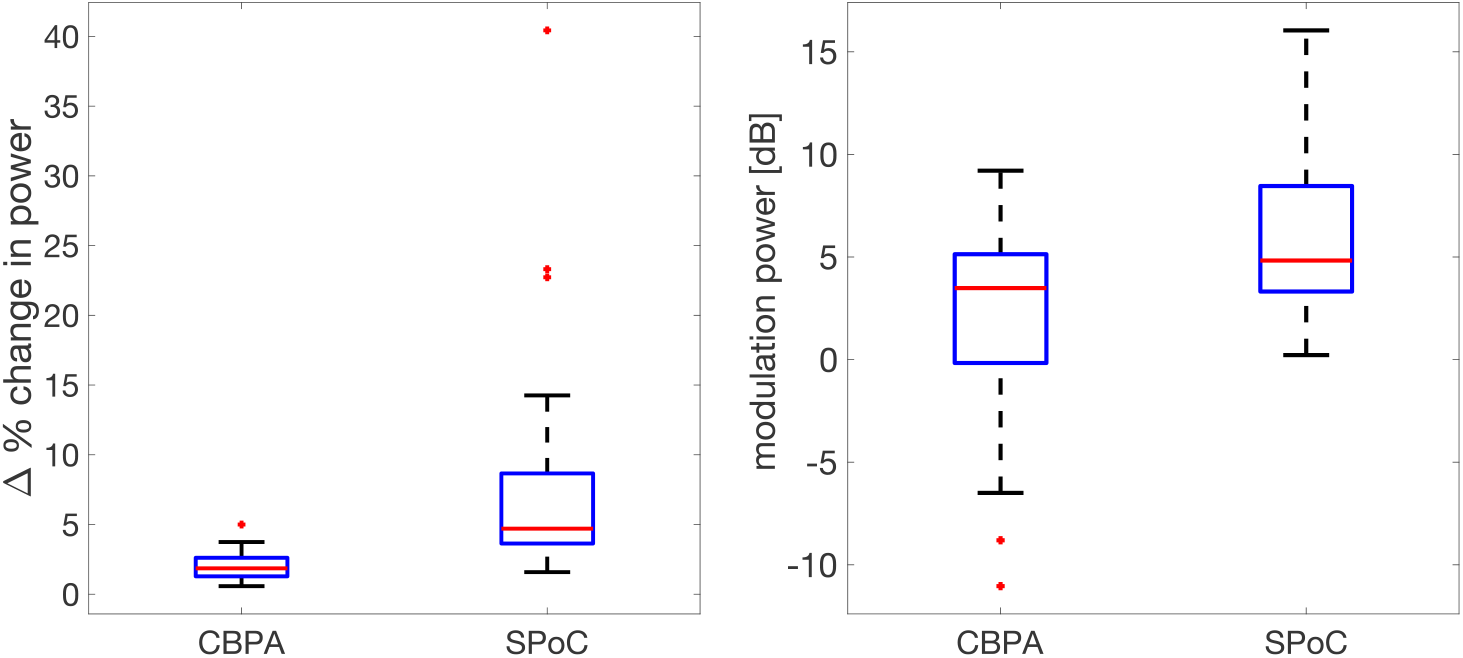
Comparative quantification of modulation at the beat frequency for CBPA and SPoC as boxplots. *Left:* Peak-to-peak difference between event-related synchronisation and desynchronisation (ERS/ERD) shows individual modulation depth. *Right:* Fast Fourier Transformation of the time frequency response at the beat frequency shows individual modulation power. Boxes show upper and lower quartiles, red line indicates the median.

We measure modulation power by applying a Fast Fourier Transform to each participant’s TFR of the first SPoC component in 8-20 Hz (Fig.5). After averaging the result over 8-20 Hz, we calculated local peaks in the FFT of the beat frequency (2.5 Hz) as the mean difference in power to its two neighboring sample points. Those “local peaks” give us a measurement for modulation power, i.e. a higher local peak means relatively more modulation power at the beat frequency. We also calculate local peaks for the TFR of the 78 significant sensors of the CBPA analysis in Fig. 1 and average over 6-18 Hz. Results are shown in Fig. 6 (*right*) as boxplots where one values is assigned per participant for SpoC and CBPA. Again, local peaks for SpoC are significantly higher (*p* = 0.0024) than for CBPA which means that TFR for SPoC shows significant higher relative modulation power in the beat frequency.

## 5 Discussion

This study aims to detect periodic modulations in oscillatory power in a pre-defined frequency band of interest in the MEG activity of participants presented with a regular, acoustic beat. We designed a customized machine learning-based analysis pipeline using two spatial filtering techniques: to first seek brain activity with maximal energy in the frequency band of interest, and then in the second step extract components that display the strongest modulation at the beat frequency within the given frequency band of interest. We test this pipeline on MEG recordings from 28 participants who had been passively listening to a 5-min long regular tone sequence with a 400 ms beat period (2.5 Hz), whilst watching a silent film. We compare the results to those from a sophisticated state-of-the-art cluster-based permutation analysis (CBPA). Both analysis pipelines employ Morlet wavelet-based time frequency analyses to obtain the spectrogram of the activity of the relevant sources and to then apply a fast-Fourier transformation (FFT) in order to compare the modulation spectra, in terms of specificity and modulation power at the beat period within the frequency band of interest. The whole pipeline calculated for two beat periods epochs that guarantee a long enough time window to most clearly see the repetitive nature of the hypothesised modulation. We show here the results for beat-related activity in two frequency bands of interest, that is the beta and the alpha band: an informed choice based on previous reports by [22] and [23] for the beta band, plus [36–38] for the alpha band.

### Outcome of f-SPoC in comparison to CBPA pipeline

Our dedicated pipeline tackles the hypothesized oscillatory power modulations by first employing Spatio-Spectral Decomposition (SSD) [1, 39] to find components with high signal power in the frequency band of interest. Secondly, we apply our customised version of the Source Power Co-modulation (SPoC) algorithm, the original version of which was published by Dähne and co-workers in 2014 [2]. The present version is Fourier-based, hence called f-SPoC, in that is extracts components in theoretical source space that modulate their activity with the period of the beat frequency, in other words that display one peak and one trough per beat period. The goal here was to find in a data-driven fashion the most relevant components that show a beat-based modulation in their spectrogram within the frequency band of interest. Following the identification of the most relevant components (here, two per participant), time-frequency analysis (TFA) was applied using Morlet wavelets [40] to obtain the spectrogram. The expected patterns indeed came out very strong, in that across both the lower beta and the alpha band they display one clear peak (event-related synchronisation, ERS) and one clear trough (event-related desynchronisation, ERD) per beat period, and that at the scale of ±5% change in power (see Figure 5).

In the comparison pipeline, we opted for a sophisticated cluster-based permutation analysis (CBPA) [3] that, like with our f-SPoC pipeline, would also be able to tackle efficiently the activity within the frequency bands of interest. We here averaged across all the sensors that showed significant activity within the frequency band of interest at the group level (78 out of 102 sensors, see Figure 1) to then perform the TFA. Also here, the spectrogram displays a repeating pattern comprising one peak (ERS) and one trough (ERD) per beat period within our frequency band of interest, discernible by eye between 6 and 18 Hz. The pattern resembles that of the f-SPoC outcome, but appears much weaker.

The modulation spectra obtained after the common final step of applying the FFT on the TFRs show the natural decrease of modulation power with increasing modulation frequency and with increasing frequency of brain (MEG) activity. In order to quantify the sought-after modulation at the beat frequency of 2.5 Hz, we calculated the “local peak” values (the mean difference to the neighbouring frequencies in power) in the FFT and show a significant gain for the SPoC compared to the CBPA-based results. In addition, our measure of modulation depth, calculated as the peak-to-trough difference in the spectrograms, was also significantly higher for the SPoC vs. the CBPA-based results.

In sum and in line with our expectation, the hypothesis-based, data-driven use of our f-SPoC pipeline serves to most efficiently seek the hypothesized beat-based power modulations. It supports the extraction of the maximally relevant theoretical source-space component activity, without anatomical priors or even anatomical information at the individual level required as an input. This renders our method versatile and applicable to both MEG and EEG time series analysis, especially when details structural data (MRI) are not available.

### Beat-based modulations in oscillatory power

Building on pre-existing work, we here sought beat-based entrainment of activity in the beta band, more specifically in low (15-20 Hz) and high (20-25 Hz) beta, as well as in the alpha band (8-12 Hz). This choice is to a certain degree related to the widely assumed, albeit not uncontroversial “typical” roles of those two frequency bands, which have recently been observed to behave in a co-modulated fashion (e.g. [41]).

Traditionally, alpha band power is associated with relaxation and has been shown to be suppressed by for instance sensorimotor processes [42]. Beta band power in turn is generally associated with states of alertness, including functions such as monitoring the status quo [43]. With relevance to timing, beta band activity has been promoted repeatedly in sensory-motor tasks, in association with anticipatory and predictive processes, especially in rhythmic contexts [44–46]. More musically oriented studies have highlighted beta band power modulations in the context of metrical beat processing, which involves two or more hierarchically organised levels of periodicity [21, 47, 48].

Two key sets of findings in motivating our present study were those presented by Fujioka et al. [22] (MEG) and Chang et al. [23] (EEG). Both show clear periodic power modulations in the beta band that entrain with a plain regular beat comprising one level of periodicity similar to our stimulus. Consistent with those studies, also we found clear modulations in the lower beta band (15-20 Hz). However, they were much less discernible in the higher beta band (20-25 Hz), where Fujioka et al. (2012) had found the strongest modulation. Following up on an earlier report by [36], we in addition looked for related periodic activity in the alpha band. Indeed, our results showed even clearer patterns when targeting the combination of alpha and lower beta together. The analysis across the two frequency bands yielded the most prominent periodic patterns of modulation in the spectrograms for both the f-SPoC and CBPA pipeline. Consistent with this, Morillon and Baillet (2017) reported phase-amplitude coupling in the power of beta as well as alpha oscillations with the period of the rate of stimulation in the delta range during passive listening [49]. In any event, the across-frequency band results for beta and alpha combined are further in line with a recent related report by Griffiths et al. (2019), who also showed an effect spanning both frequency bands, although at a slower stimulation rate [38]. Directly relevant, most recent MEG work by Lerousseau and co-workers [50] presents with yet differing results with respect to the oscillatory modulations in the relevant frequency bands in question. In their study of passive listening to a regular beat in the relevant range, in fact of identical tempo to ours (2.5 Hz), the authors found no beat-based modulation in the beta nor in the alpha band, but rather an overall suppression of activity across both bands in the auditory cortex. Some of the apparent inconsistencies between findings on the presence of beat-based modulation, in terms of the degree as well the timing of it, may remain inconciliable whilst some may be due to differences in analysis details and anatomical priors, which we briefly address in the following two sections.

### Evoked vs. induced activity

The separation of “evoked” from “induced” activity has become an influential idea over the past decades and aims to distinguish phase-locked neural processes, which are invariant in their timing to a given stimulus from those that vary in their exact timing from trial to trial, i.e. typically oscillations in the higher frequency bands. In the context of timing, for instance, the idea might be to possibly disentangle the concrete processing of a given stimulus (e.g. a tone) from that of the abstract rhythmic entrainment, whereby the latter might be expected to serve the process underlying the prediction of the next tone in a sequence and our “feeling of the beat”.

The report on beat-based modulations by Fujioka et al. (2012) in the oscillatory beta-band power indeed focused their analysis on induced activity, i.e. on that part of the neural response that remains after subtracting the average event-related response before then calculating the time frequency response (spectrogram). A number of recent reports in contrast presented results based on total power instead, as we do here. Morillon and Baillet [49] for instance, reported phase-amplitude coupled theta-beta oscillations, in other words, periodic bursts of MEG beta band activity in total power that followed the timing of a regular acoustic beat. In another relevant recent pre-print report by Balasubramaniam’s group on an EEG study on an audio-visual paradigm involving a regular rhythm [51], the authors have also not been able to reliably show beat-based modulations of beta power in induced activity - but in evoked activity instead. Similarly, and in line with what we observe here, in total but not in induced activity, Tavano et al. (2019) presented rhythmic modulation of total power within the lower beta band (15-19 Hz, or “Beta 1”, [52]). The authors demonstrated a peak at about 100 ms before the next tone in a regular sequence played at 4 Hz. There were no clear effects in alpha power, which may, as the authors discuss, have to do with the paradigm using an active listening task focussing on target tones. More specifically, the task included an informed and an uninformed condition, which did have an effect on the strength of the beta band modulation in the study by Tavano et al. (2019). For the interpretation of our results, this might suggest an underestimation of the room for strength of modulation in the absence of a task, i.e. that modulation might be stronger when using an active task.

Related observations in terms of total or evoked vs. induced power have been made in the alpha band. Consistent with this are other reports of phase-locked alpha activity by for instance Escoffier et al. (2015) on the effects of auditory rhythmic stimulation on visually evoked oscillations and event-related potentials [53]. One relevant study by Arnal, Doelling & Poeppel (2015) [54] reported a periodic power modulation, i.e. a phase-amplitude coupling with the “delta” rhythm of auditory stimulation, in the beta band as well as in the alpha band. The authors do not explicitly discuss a differentiation of induced vs. evoked activity, and we assume as with the majority of studies discussed in this section, that also they are looking at total power. The neural process underlying this phenomenon of “evoked oscillations” might be a phase resetting mechanism, as described before for the alpha band [55], as well as even for the gamma band [56].

### Systems neuroscience considerations

The data-driven nature of our pipeline comes with no need for anatomical priors, neither in general nor at the individual level. The SPoC-based extraction of components is thus in no way specific limited to directing it toward any particular anatomical structure. An extracted component might in theory reflect the activity of one single brain structure, or just as likely that between two or more.

In the first instance, we expected to pick up a component that would reflect the induced beta-band power oscillations in the auditory cortex as reported by Fujioka and colleagues [22]. However, our analysis did not yield a pattern that resembled those. Curiously, the periodic modulations that we did find in our analysis looking at total power present with a slight shift in the timing of the observed peak and troughs, i.e. in their latency with respect to the beat, in comparison to Fujioka et al.’s (2012) results. This difference could in part be due to differences in stimulation, i.e. a continuous 5-min sequence with one tempo in our case, this might also be due to a difference in the source(s) we pick up in our components.

The relevant SPoC components displaying a beat-based modulation in their oscillatory power that we extract here may well reflect the communication between two or more brain structures as proposed by Fries (2005) [57] and Lakatos et al. (2019) [58]. This could for instance be a projection between auditory areas in the temporal lobe and motor areas of the frontal lobe, or might even include a subcortical component. Such a network activity would be consistent with previous observations of beta being related to timing network processes, i.e. in the form of projections from motor cortex to auditory, as elaborated by Khanna and Carmena, 2019 [59], or the beta phase synchronization in the frontal-temporal-cerebellar network during auditory-to-motor rhythm learning [60]. Another related recent MEG study of rhythmic entrainment and mismatch detection in the order of predictable sounds sequences, demonstrated a differential effect in the phase-amplitude coupling of the alpha band in the feed-forward vs. the feed-back projection between the hippocampus, auditory cortex and frontal areas [61]. Along these lines, an elegant MEG study in a cohort of patients with non-fluent aphasia due to frontal neurodegeneration compared to a healthy control group employed acoustic degradation of speech stimuli and was able to distinguish causal top-down beta band signals from frontal speech temporal auditory brain regions, carried in the beta band [62].

The current approach however does not try to specifically address or distinguish between any or all of the above possibilities. It is thereby fundamentally different to any pre-informed source-analysis methods (discrete and continuous) that are ideally used in combination with a brain scan (MRI) or prototype brain model. The aim here is to most efficiently and precisely extract the relevant oscillatory activity within a given band of interest in theoretical source space by a customised, data-based ML-based analysis, without the need for anatomical priors or limitations in confining to pre-defined regions of interest. In theory, one might expect our SPoC pipeline to pick up on two or more relevant separable components, that may even correspond to different brain structures or networks. However, that was not the case here, and instead we find one prominent main component that displays beat-based modulation of oscillations across both the alpha and lower beta band.

### Functional interpretations

Whilst the aim of one comprehensive, unifying interpretation of the data available may currently appear unattainable, one shared observation made by a number of the studies discussed above is that of time-locked modulations in both beta and alpha power in the context of a regular rhythm. Within both these frequency bands, the converging evidence shows one peak per periodically ocurring sensory and motor event, whereby the exact latency of the peak varies between studies, possibly to do with differences in paradigms, as does the proportion of evoked and induced activity reported. The functional roles and criticalness of this modulation remains debatable and the different studies have different emphases on motor or sensory processes and their interaction. However, they generally point into the directions of temporal expectation and predictive processing, with a possible differentiation between anticipating and monitoring of sensory stimuli or movement preparation.

Arnal et al. (2015) for instance, in their study on delay detection in rhythmic tone sequences, interpreted differentially the beta band peak as a means of temporal prediction that precedes an upcoming stimulus, and the alpha peak to be used by listeners to “update” their prediction afterward [54]. Of note, the authors used a relatively slow tempo (800-1200 ms) compared to ours and most of the other relevant beat-based studies typically employing a beat rate of about 2 Hz (500 ms), or a range of inter-onset-intervals of up to about 800 ms in case of non-isochronous rhythm. Whilst this relatively slow tempo may in part tap into processes of supra-second timing and its categorial distinction from sub-second timing made elsewhere [63], it might be what aided the distinction between the two peaks and their functional interpretation.

A number of other studies have emphasised the interaction between sensory predictive, processing of rhythmic stimulation with activity in the motor system. Morillon and Baillet (2017; MEG) for instance documented interaction between auditory regions and sensorimotor cortex in the form of interdependent oscillations in the beta band, and in addition a trend in the alpha band, with the beat frequency (1.5 Hz), in a temporal prediction task based on two separable acoustic streams, and found an increase in activity by adding rhythmic movement [49]. Along these lines, Edagawa and Kawasaki (2017) showed an increase in the beta power and its phase synchronization within the frontal-temporal-cerebellar network during rhythmic auditory-to-motor learning in a delayed reproduction task [60]. Other authors promote even more explicitly the use of beta oscillations of the motor system for sensory predictions [44] in addition to their likely primary function of the modulation of neural excitability for in movement preparation. A related recent study combining somatosensory stimulation with TMS applied over the motor cortex strongly supports the role for sensorimotor interaction in the genesis of a beta rebound peak [64]. This is consistent with the proposed role for beta oscillations of the motor system for sensory predictions [44], in addition to its likely primary function in timed movement preparation [65, 66] and further supported by the rebound peak latency scaling with the period in a rhythmic motor task [65]. In terms of a correlation with behaviour in the sensory domain, as suggested for rhythmic motor learning by [60], or a possible specific role in the processing of speech [67, 68], further work is needed.

## 6 Conclusion

This work presents a new approach to finding beat-based entrainment in the human brain, that is data-driven and geared to detect source-space components’ activity that maximally corresponds to a given target function. More specifically, we employed a customised version of the machine learning-based source power co-modulation algorithm (SPoC) to extract source space component activity that maximally entrains its power within the frequency band of interest with the given beat frequency, based on Fourier analysis (hence, f-SPoC). Optimal results could be obtained by first applying SSD as a pre-processing step to maximize signal power in the band of interest (here, lower beta plus alpha combined) and feeding the SSD components with the highest information content (here, fifteen) into the f-SPoC algorithm in order to extract those source component/s that show a clear modulation of power at the beat frequency (here, Hz). The f-SPoC pipeline clearly outperformed our sophisticated sensor-space comparison analysis based on a state-of-the-art cluster-based permutation technique. Future applications will include but are not limited to its use in combination with active sensing tasks, more musically oriented or speech-based paradigms, its implementation in the EEG, and in working with those with neurological or developmental disorders of rhythm.

## Acknowledgements

We thank all the participants for taking part in the study. This work was partly funded by the German Ministry for Education and Research as BIFOLD - Berlin Institute for the Foundations of Learning and Data (ref. 01IS18025A and ref 01IS18037A), awarded to SB. Financial support also came from the Danish National Research Foundation (project number DNRF117 awarded to PV) and from the People Programme (Marie Curie Actions) of the European Union’s Seventh Framework Programme (FP7/2007-2013) under REA grant agreement no. 600209 (TU Berlin/IPODI) awarded to MG. We further thank our colleagues Angus Stevner for advice on pre-processing and Alexander Kraut for numerous hours of showing the way into MEG acquisition. Huge thanks go to Anastasiia Popova and Antonieta Martínez-Guerrero for continuous and tireless work over the two months of data acquisition. The authors remain grateful to Klaus-Robert Müller, Chair of Machine Learning Group at TU Berlin, for his continued support, including his input and hosting of the initial idea to develop the present customised version of SPoC in order to track beat-based modulation of activity in the human brain.

## References

1. Nikulin VV, Nolte G, Curio G. A novel method for reliable and fast extraction of neuronal EEG/MEG oscillations on the basis of spatio-spectral decomposition. NeuroImage. 2011;55(4):1528–1535.

2. Dähne S, Meinecke FC, Haufe S, Höhne J, Tangermann M, Müller KR, et al. SPoC: a novel framework for relating the amplitude of neuronal oscillations to behaviorally relevant parameters. NeuroImage. 2014;86:111–122.

3. Maris E, Oostenveld R. Nonparametric statistical testing of EEG-and MEG-data. Journal of neuroscience methods. 2007;164(1):177–190.

4. London J, Time HI. Psychological Aspects of Musical Meter; 2004.

5. Bekius A, Cope TE, Grube M. The beat to read: a cross-lingual link between rhythmic regularity perception and reading skill. Frontiers in human neuroscience. 2016;10:425.

6. Grahn JA, Brett M. Impairment of beat-based rhythm discrimination in Parkinson’s disease. Cortex. 2009;45(1):54–61.

7. Huss M, Verney JP, Fosker T, Mead N, Goswami U. Music, rhythm, rise time perception and developmental dyslexia: perception of musical meter predicts reading and phonology. Cortex. 2011;47(6):674–689.

8. Ladányi E, Persici V, Fiveash A, Tillmann B, Gordon RL. Is atypical rhythm a risk factor for developmental speech and language disorders? Wiley Interdisciplinary Reviews: Cognitive Science. 2020; p. e1528.

9. Bengtsson SL, Ullen F, Ehrsson HH, Hashimoto T, Kito T, Naito E, et al. Listening to rhythms activates motor and premotor cortices. cortex. 2009;45(1):62–71.

10. Chen JL, Penhune VB, Zatorre RJ. Listening to musical rhythms recruits motor regions of the brain. Cerebral cortex. 2008;18(12):2844–2854.

11. Grahn JA, Brett M. Rhythm and beat perception in motor areas of the brain. Journal of cognitive neuroscience. 2007;19(5):893–906.

12. Ivry RB, Keele SW. Timing functions of the cerebellum. Journal of cognitive neuroscience. 1989;1(2):136–152.

13. Grube M, Lee KH, Griffiths TD, Barker AT, Woodruff PW. Transcranial magnetic theta-burst stimulation of the human cerebellum distinguishes absolute, duration-based from relative, beat-based perception of subsecond time intervals. Frontiers in Psychology. 2010;1:171.

14. Teki S, Grube M, Kumar S, Griffiths TD. Distinct neural substrates of duration-based and beat-based auditory timing. Journal of Neuroscience. 2011;31(10):3805–3812.

15. Grahn JA, Rowe JB. Finding and feeling the musical beat: striatal dissociations between detection and prediction of regularity. Cerebral cortex. 2013;23(4):913–921.

16. Cope TE, Grube M, Singh B, Burn DJ, Griffiths TD. The basal ganglia in perceptual timing: timing performance in Multiple System Atrophy and Huntington’s disease. Neuropsychologia. 2014;52:73–81.

17. Grube M, Cooper FE, Chinnery PF, Griffiths TD. Dissociation of duration-based and beat-based auditory timing in cerebellar degeneration. Proceedings of the National Academy of Sciences. 2010;107(25):11597–11601.

18. Teki S, Grube M, Griffiths TD. A unified model of time perception accounts for duration-based and beat-based timing mechanisms. Frontiers in integrative neuroscience. 2012;5:90.

19. Toiviainen P, Burunat I, Brattico E, Vuust P, Alluri V. The chronnectome of musical beat. NeuroImage. 2020;216:116191.

20. Large EW, Herrera JA, Velasco MJ. Neural networks for beat perception in musical rhythm. Frontiers in systems neuroscience. 2015;9:159.

21. Iversen J, Repp B, Patel A. Top-down control of rhythm perception modulates early auditory responses. Annals of the New York Academy of Sciences. 2009;1169(1):58–73.

22. Fujioka T, Trainor LJ, Large EW, Ross B. Internalized timing of isochronous sounds is represented in neuromagnetic beta oscillations. Journal of Neuroscience. 2012;32(5):1791–1802.

23. Chang A, Bosnyak DJ, Trainor LJ. Unpredicted pitch modulates beta oscillatory power during rhythmic entrainment to a tone sequence. Frontiers in psychology. 2016;7:327.

24. Teki S. Beta drives brain beats. Frontiers in Systems Neuroscience. 2014;8:155.

25. Fujioka T, Trainor L, Large E, Ross B. Beta and gamma rhythms in human auditory cortex during musical beat processing. Annals of the New York Academy of Sciences. 2009;1169(1):89–92.

26. Association WM, et al. World Medical Association Declaration of Helsinki. Ethical principles for medical research involving human subjects. Bulletin of the World Health Organization. 2001;79(4):373.

27. Taulu S, Simola J. Spatiotemporal signal space separation method for rejecting nearby interference in MEG measurements. Physics in Medicine & Biology. 2006;51(7):1759.

28. Taulu S, Hari R. Removal of magnetoencephalographic artifacts with temporal signal-space separation: demonstration with single-trial auditory-evoked responses. Human brain mapping. 2009;30(5):1524–1534.

29. Oostenveld R, Fries P, Maris E, Schoffelen JM. FieldTrip: open source software for advanced analysis of MEG, EEG, and invasive electrophysiological data. Computational intelligence and neuroscience. 2011;2011.

30. Makeig S, Bell AJ, Jung TP, Sejnowski TJ. Independent component analysis of electroencephalographic data. In: Advances in neural information processing systems; 1996. p. 145–151.

31. Basu D. Randomization analysis of experimental data: the Fisher randomization test. In: Selected Works of Debabrata Basu. Springer; 2011. p. 305–325.

32. Sinkkonen J, Tiitinen H, Näätänen R. Gabor filters: an informative way for analysing event-related brain activity. Journal of neuroscience methods. 1995;56(1):99–104.

33. Pfurtscheller G, Da Silva FL. Event-related EEG/MEG synchronization and desynchronization: basic principles. Clinical neurophysiology. 1999;110(11):1842–1857.

34. Haufe S, Meinecke F, Görgen K, Dähne S, Haynes JD, Blankertz B, et al. On the interpretation of weight vectors of linear models in multivariate neuroimaging. Neuroimage. 2014;87:96–110.

35. Cirelli LK, Bosnyak D, Manning FC, Spinelli C, Marie C, Fujioka T, et al. Beat-induced fluctuations in auditory cortical beta-band activity: using EEG to measure age-related changes. Frontiers in psychology. 2014;5:742.

36. Fujioka T, Ross B. Auditory processing indexed by stimulus-induced alpha desynchronization in children. International journal of psychophysiology. 2008;68(2):130–140.

37. Rohenkohl G, Nobre AC. Alpha oscillations related to anticipatory attention follow temporal expectations. Journal of Neuroscience. 2011;31(40):14076–14084.

38. Griffiths BJ, Mayhew SD, Mullinger KJ, Jorge J, Charest I, Wimber M, et al. Alpha/beta power decreases track the fidelity of stimulus-specific information. Elife. 2019;8:e49562.

39. Haufe S, Dähne S, Nikulin VV. Dimensionality reduction for the analysis of brain oscillations. NeuroImage. 2014;101:583–597.

40. Cohen MX. Analyzing neural time series data: theory and practice. MIT press; 2014.

41. Bauer M, Stenner MP, Friston KJ, Dolan RJ. Attentional modulation of alpha/beta and gamma oscillations reflect functionally distinct processes. Journal of Neuroscience. 2014;34(48):16117–16125.

42. Arroyo S, Lesser RP, Gordon B, Uematsu S, Jackson D, Webber R. Functional significance of the mu rhythm of human cortex: an electrophysiologic study with subdural electrodes. Electroencephalography and clinical Neurophysiology. 1993;87(3):76–87.

43. Engel AK, Fries P. Beta-band oscillations—signalling the status quo? Current opinion in neurobiology. 2010;20(2):156–165.

44. Arnal LH. Predicting “when” using the motor system’s beta-band oscillations. Frontiers in human neuroscience. 2012;6:225.

45. Arnal LH, Giraud AL. Cortical oscillations and sensory predictions. Trends in cognitive sciences. 2012;16(7):390–398.

46. Gupta DS, Chen L. Brain oscillations in perception, timing and action. Current Opinion in Behavioral Sciences. 2016;8:161–166.

47. Fujioka T, Zendel BR, Ross B. Endogenous neuromagnetic activity for mental hierarchy of timing. Journal of Neuroscience. 2010;30(9):3458–3466.

48. Fujioka T, Ross B, Trainor LJ. Beta-band oscillations represent auditory beat and its metrical hierarchy in perception and imagery. Journal of Neuroscience. 2015;35(45):15187–15198.

49. Morillon B, Baillet S. Motor origin of temporal predictions in auditory attention. Proceedings of the National Academy of Sciences. 2017;114(42):E8913–E8921.

50. Lerousseau JP, Trébuchon A, Morillon B, Schön D. Persistent neural entrainment in the human cortex is frequency selective. bioRxiv. 2019; p. 834226.

51. Comstock DC, Ross JM, Balasubramaniam R. Modality-specific frequency band activity during neural entrainment to auditory and visual rhythms. bioRxiv. 2020;.

52. Tavano A, Schröger E, Kotz SA. Beta power encodes contextual estimates of temporal event probability in the human brain. PLoS ONE. 2019;14(9):e0222420.

53. Escoffier N, Herrmann CS, Schirmer A. Auditory rhythms entrain visual processes in the human brain: Evidence from evoked oscillations and event-related potentials. NeuroImage. 2015;111:267–276.

54. Arnal LH, Doelling KB, Poeppel D. Delta–beta coupled oscillations underlie temporal prediction accuracy. Cerebral Cortex. 2015;25(9):3077–3085.

55. Min BK, Busch NA, Debener S, Kranczioch C, Hanslmayr S, Engel AK, et al. The best of both worlds: phase-reset of human EEG alpha activity and additive power contribute to ERP generation. International Journal of Psychophysiology. 2007;65(1):58–68.

56. Shamir M, Ghitza O, Epstein S, Kopell N. Representation of time-varying stimuli by a network exhibiting oscillations on a faster time scale. PLoS Comput Biol. 2009;5(5):e1000370.

57. Fries P. A mechanism for cognitive dynamics: neuronal communication through neuronal coherence. Trends in cognitive sciences. 2005;9(10):474–480.

58. Lakatos P, Gross J, Thut G. A new unifying account of the roles of neuronal entrainment. Current Biology. 2019;29(18):R890–R905.

59. Khanna P, Carmena JM. Neural oscillations: beta band activity across motor networks. Current opinion in neurobiology. 2015;32:60–67.

60. Edagawa K, Kawasaki M. Beta phase synchronization in the frontal-temporal-cerebellar network during auditory-to-motor rhythm learning. Scientific reports. 2017;7(1):1–9.

61. Recasens M, Gross J, Uhlhaas PJ. Low-frequency oscillatory correlates of auditory predictive processing in cortical-subcortical networks: a MEG-study. Scientific reports. 2018;8(1):1–12.

62. Cope TE, Sohoglu E, Sedley W, Patterson K, Jones P, Wiggins J, et al. Evidence for causal top-down frontal contributions to predictive processes in speech perception. Nature Communications. 2017;8(1):1–16.

63. Lewis PA, Miall RC. Brain activation patterns during measurement of sub-and supra-second intervals. Neuropsychologia. 2003;41(12):1583–1592.

64. Cardellicchio P, Hilt PM, Dolfini E, Fadiga L, D’Ausilio A. Beta rebound as an index of temporal integration of somatosensory and motor signals. Frontiers in Systems Neuroscience. 2020;14.

65. Salmelin R, Hámáaláinen M, Kajola M, Hari R. Functional segregation of movement-related rhythmic activity in the human brain. Neuroimage. 1995;2(4):237–243.

66. Kilavik BE, Zaepffel M, Brovelli A, MacKay WA, Riehle A. The ups and downs of beta oscillations in sensorimotor cortex. Experimental neurology. 2013;245:15–26.

67. Gross J, Hoogenboom N, Thut G, Schyns P, Panzeri S, Belin P, et al. Speech rhythms and multiplexed oscillatory sensory coding in the human brain. PLoS Biol. 2013;11(12):e1001752.

68. van Bree S, Sohoglu E, Davis MH, Zoefel B. Sustained neural rhythms reveal endogenous oscillations supporting speech perception. PLoS Biol. 2021;19(2):e3001142.

